# Validation of binary non-targeted methods: mathematical framework and experimental designs

**DOI:** 10.1101/2021.01.19.427235

**Authors:** Steffen Uhlig, Kapil Nichani, Manfred Stoyke, Petra Gowik

## Abstract

Non-targeted methods (NTMs) are being increasingly developed and adopted to detect food fraud and identify the authenticity of the food. Method validation is a critical step before bringing the NTMs to the routine to be assured of method performance and trust that its outcomes are reliable. However, the paucity of well structured and harmonized validation strategies has been one of the hurdles, withholding the exploitation of NTMs. This report aims to describe a validation framework for methods that involve binary classification, which are prevalent in non-targeted workflows. We foresee this work to contribute to filling the current gap in the provisions for NTM method validation; perhaps also push the dialogue further to collectively resolve existing challenges.

## 1. Introduction

Food fraud incidences remain pervasive, requiring diverse monitoring and inspection mechanisms along with the food production, processing, and supply chain [1,2]. Analytical measurement methods play a central role in investigating the authenticity of food, detecting adulteration or mixing, or determining geographical origin [3–5]. Recent advancements in instrument technologies, boasting superior speed and resolution, along with a reduction in the cost of analysis, have led to the manifold adoption of non-targeted methods (NTMs) [6,7].

It is unsurprising that NTMs are not limited to measuring predetermined analytes of interest through its suggestive name [8]. Instead, the measurement process yields spectra or genomic sequences that form characteristic “fingerprints” for the sample. Spectra can be the result of methods involving chromatographic separation followed by mass spectrometry measurements [9], or direct spectral methods such as nuclear magnetic resonance (NMR) [10], Fourier transformed infrared (FTIR) [11], or Raman spectroscopy [12]. On the other hand, sequences can be acquired through massively parallel sequencing of shotgun or amplicon sequences, sequenced using next-generation sequencing technologies [13,14]. A large assemblage of published work chronicles the assortment of methods, describing the salient features of different NTMs applied to diverse food types or objectives around detecting food fraud. For a compilation of such approaches, the reader is referred to literature elsewhere [3,6,15–17].

A large subset of NTMs can be formulated as a binary classification problem [18,19]. For instance – is the meat adulterated [20]? Or has the basmati rice been sourced from a particular region [21]? Or is honey mixed with sugar syrup [10]? Hence, it is not only appropriate to focus on such NTMs, but also consequential because a large number of methods are qualitative in nature.

A prerequisite to using any analytical method for testing samples on a routine basis is that the method should be validated, whereby its performance characteristics should be defined and suitably assessed [22]. In order to match the pace of new methods being proposed and developed, there is a need to devise new concepts for method validation and establish them for the benefit of producers, consumers, and regulators alike [15]. A validation procedure, in general, involves determining the risk of a false positive or false negative. For the method developer, this is important to demonstrate the applicability for intended use objectively; for the method user, it is crucial for quality assurance and accreditation [23].

To this end, this work describes a basis for a validation strategy and evaluation of performance characteristics. This report is structured in the following way. First, it describes a general mathematical formulation of a binary NTM problem. Then it suggests experimental designs in order to realize a method validation study. Herein, considerations for the minimum number of samples are also discussed. Lastly, it proposes a statistical model for evaluating pertinent performance characteristics.

## 2. Mathematical Framework

We assume that the samples *i* to be analyzed are from one of two classes (sets), A or B (*i* ∈ *A* or *i* ∈ *B*). The measurement outcome is either a spectrum or a sequence. This result can be expressed as a one, two, or multi-dimensional matrix of real-valued numbers. We denote this matrix by *X*_*i*_. It is the basis of the pipeline y, which produces a result 0 or 1 for class A or B, respectively, based on an NTM, as seen in Equation 1.

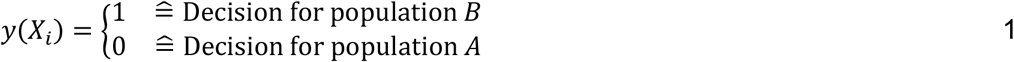

For each measurement *X*_*i*_ of sample *i* from one of the two classes B or A, a result can be either correct or incorrect. Table 1 describes the possible outcomes.

**Table 1:** Contingency table, showing the different outcomes for measuring i samples that can belong either to classes A or B, using a pipeline y, resulting in a matrix of values *X*_*i*_.

| | $i \in A$ | $i \in B$ |
| --- | --- | --- |
| $y(X_i) = 0$ | True Negative (TN) | False Negative (FN) |
| $y(X_i) = 1$ | False Positive (FP) | True Positive (TP) |

Here it should be noted that the measurement *X*_*i*_has a random character because repeating the measurement on the same sample could result in a different value. Consequently, depending on the sample i to be measured, a probability can be specified for each of the events – true negative (TN), true positive (TP), false negative (FN), and false-positive (FP). These are shown in Table 2. The determined probabilities for specificity(i) and sensitivity(i) depend on the sample i investigated in each case. They can be determined by repeated measurements of such a sample i, for example, within the framework of an interlaboratory test.

**Table 2:**
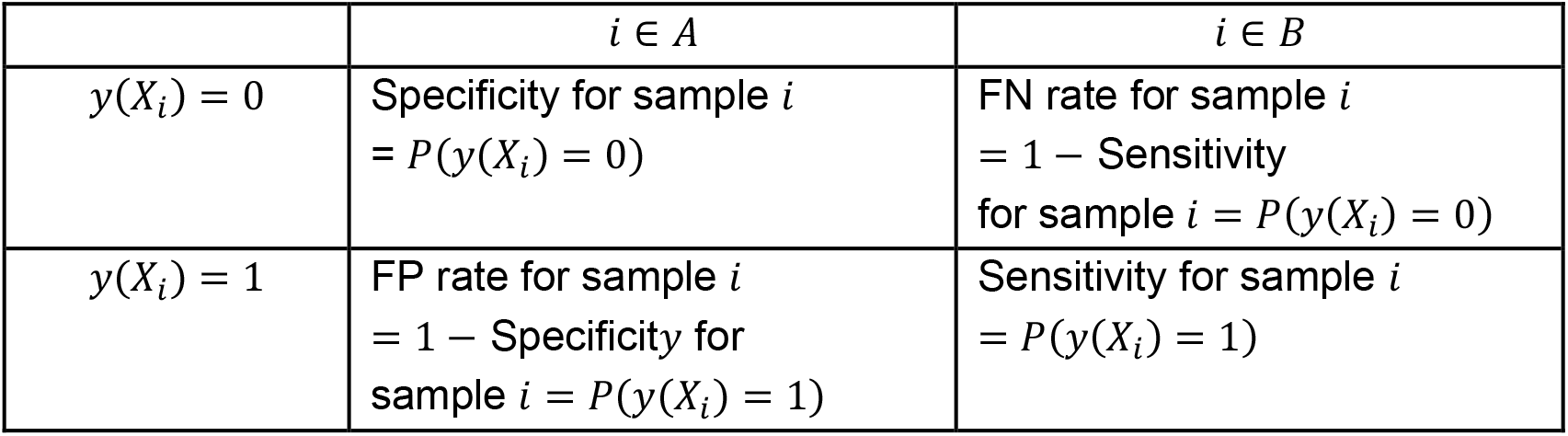
Contingency table, with the formulation for probabilities for TN, TP, FN, and FP.

**Table 3:**
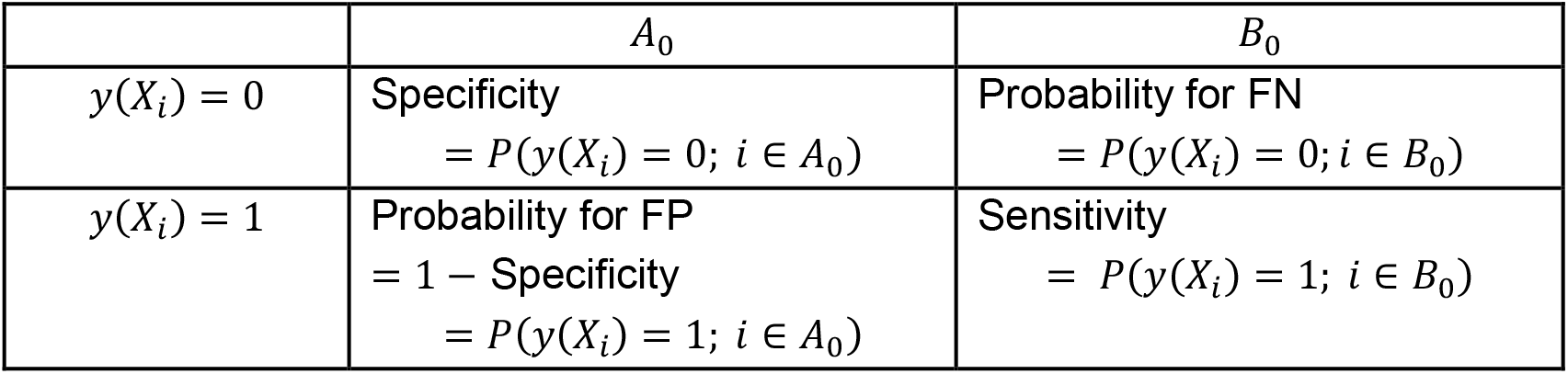
Contingency table with the formulation of the sensitivity and specificity with regard to sub classes *A*_0_ ⊆ *A* and *B*_0_ ⊆ *B*.

One often does not know very much about the probabilities, as given in Table 2. Nevertheless, it is not merely a theoretical construct. If a laboratory performs an NTM, it will obtain either a positive or a negative result. If the result is positive, the question arises as to the probability with which a false positive result has occurred: this is the specificity for sample i. However, if the result is negative, the reverse question is whether a false negative has occurred: this would be the sensitivity for sample *i*. The probabilities given in Table 2 are, therefore, the two possible uncertainties associated with an NTM for sample *i*, which should be known according to ISO 17025 [24].

In order to be able to assess the performance of the NTM as a whole, the question now arises not only as to sensitivity and specificity for a fixed sample, but also as to whether or how specificity can be determined over the entire quantity A, and sensitivity over the entire quantity B. We assume that A represents the range of goods of a certain food, which - at least with regard to a certain aspect - has been declared correctly, while B represents another part of the range of goods, which, however, has been declared incorrectly and thus cannot be distinguished from A without appropriate measurement. For example, if A denotes the totality of spelt flours offered and correctly declared in Germany, B could be, for example, the totality of wheat flours offered. In this case, it is at least theoretically possible to determine the average specificity of the NTM for class A, i.e., the average value of the specificity over all spelt flours offered. This makes it possible to answer the question of the certainty with which a correctly declared product is recognized as such. Furthermore, in this case, it is also possible to determine the average sensitivity of the NTM, i.e., the average value of the sensitivity for all incorrectly declared wheat flours. This allows us to answer the question of the certainty with which incorrectly declared goods are detected as such. The formal definition of these performance parameters is described in general form in the following table. It is assumed that, depending on the problem, classes or subclasses *A*_0_ ⊆ *A* and *B*_0_ ⊆ *B* are defined, for which specificity and sensitivity are then calculated.

It is also useful to determine and state the variability of the sample-specific sensitivity and specificity over the respective populations considered. This information is particularly important if the uncertainty of the measurement results is to be determined or if the number of repeat measurements to be performed, if any, is to be specified. As illustrated in Figure 1, consider that population B is of the adulterated samples. Samples with a greater extent of adulteration can typically be easily detected with a good enough NTM and hence have high sensitivity. Conversely, samples with a low degree of adulteration will be of low sensitivity. When there is a large separation between populations A (unadulterated) and B (e.g., greater than 10 % adulteration), then unambiguous discrimination can be achieved, with low FP and FN rates. In contrast, when the separation is not very large, higher FP and FN rates can be expected. This is often the case in real-world scenarios where adulteration is either low or intended to be unobtainable. To evaluate the performance of a classification method, it is necessary to know the average FP and FN rates and how much these probabilities can vary from sample to sample.

**Figure 1.**
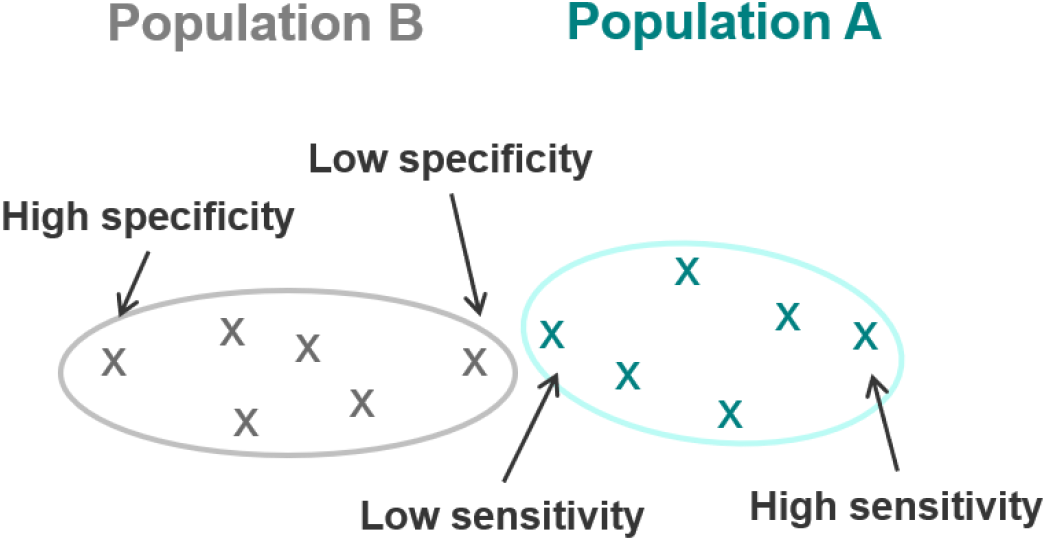
Illustration of the sample specific sensitivities and specificities. Considering population B represents adulterated samples, larger extent of adulteration will typically have high sensitivity.

## 3. Experimental designs

In order to calculate the probabilities described in the previous section, validation studies are performed. In this section, the requirements for the corresponding validation experiments are discussed.

### 3.1 Minimum requirements for the validation study

Consider an example of an NTM to evaluate the origin of the asparagus grown in Germany [6]. For the selected samples to be representative, commonly grown asparagus must be included in the validation study. It would not make sense to conduct a validation study with exotic samples originating from regions that only account for a small ratio of the total asparagus grown (e.g., Mecklenburg-Western Pomerania region growing far less asparagus than Lower Saxony, North Rhine-Westphalia, and Brandenburg). Besides, a validation study using samples limited to early harvested asparagus and not the ones sold in stores in the middle or end of the season stand questioning. Furthermore, accounting for the year of harvest is also important. Asparagus cultivated in parched years can show little resemblance in characteristics to the ones grown in a very wet year. In all, the selection of samples must be rationalized to take into account as many factors that typically contribute to differences in measurable characteristics.

The selection process of samples used for the validation study should be carried out according to statistical requirements to ensure representativeness regarding the defined population of authentic samples. Of particular importance is the consideration of the case that in the validation of the NTM, not a single false positive or a false negative sample was observed. The implications for such a case should be carefully considered, especially when the number of samples in the validation is low, i.e., only in the double digits.

Notably, the number of samples to be measured remains a contentious issue [3]. For this purpose, it can be useful to look at a subset of the total population, for example, 3% or 10%, and determine the probability that this subset is not included in a random sample. This probability is shown for a random selection of the different number of validation samples (see Figure 2).

**Figure 2.**
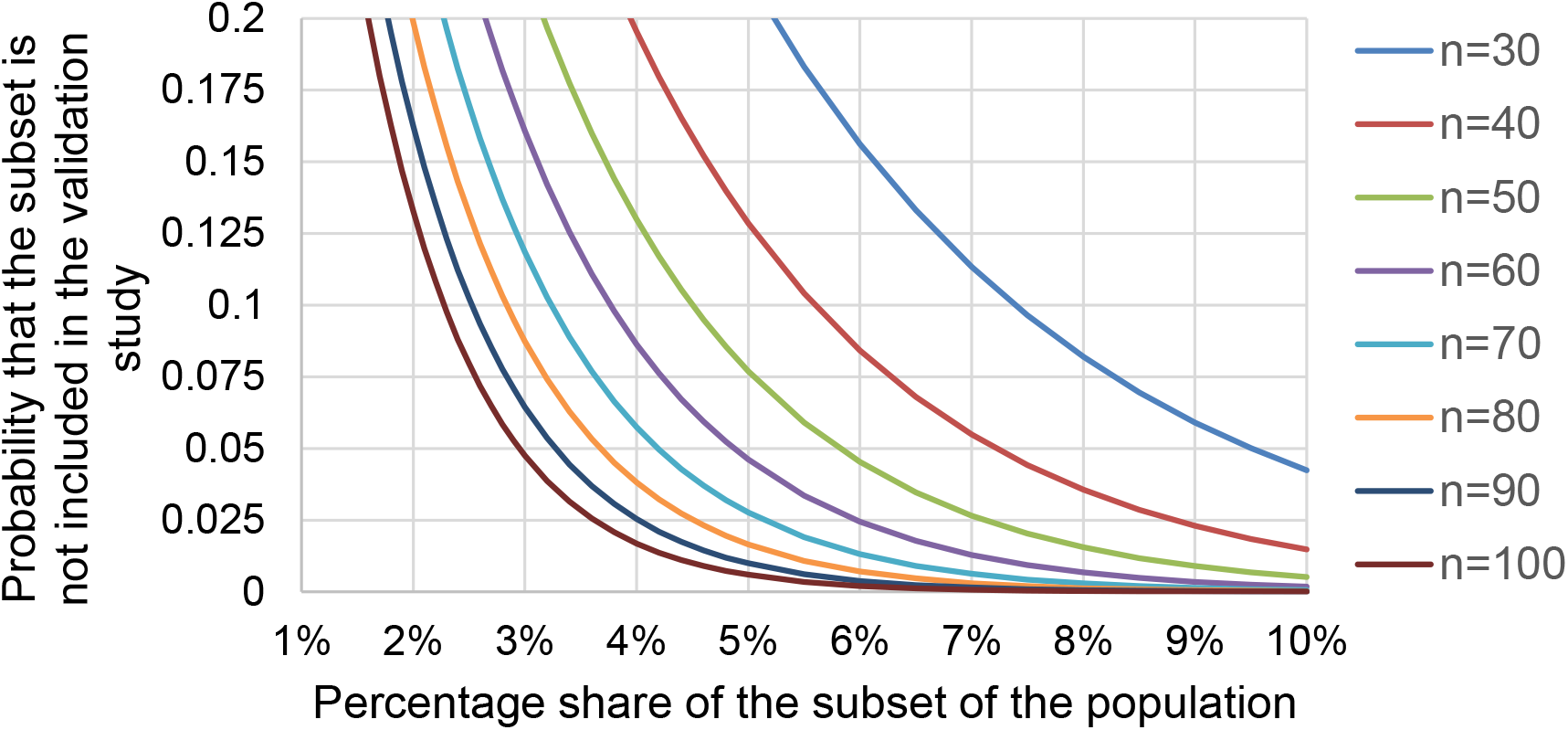
Plot showing the relation between the percentage share of the subset of the population and the probability that the subset is not included in the validation study, varying for the different number of samples (from n=30 to n=100).

As it might be expected, the probability of a particular subpopulation not being included in the validation study decreases with the size of that study. For instance, with 30 samples in the validation study, a population share of just under 10% ensures that this population share will be included in the study, with a statistical certainty of 95%. This could only be at most a few samples. If it is to be guaranteed that the samples include a subpopulation with a proportion of 5%, then at least 60 samples are required. If this requirement is set for sensitivity and specificity, 60 samples from population Q and another 60 samples from population R are needed.

### 3.2 Method validation interlaboratory study (conventional)

In order to ensure that 60 randomly selected samples from each of the two populations, A and B, are investigated, a two-step procedure is recommended. In the first step, the 60+60 samples are examined by two laboratories only. In the second step, a random selection of these samples from a total of 8 (i.e., an additional six laboratories) is then examined. Lastly, the data from both Steps 1 and 2 are evaluated.

***Step 1:*** Four laboratories perform measurements for 60 representative samples of population A and 60 representative samples of population B according to a standardized procedure with uniform sample preparation. Two laboratories analyze each sample, preferably one with a superior (or new) instrument and the other with an average (or old) instrument. The assignment of the randomly selected samples to the laboratories in an orthogonal fashion is shown in Table 4. No replicates are needed so that each laboratory conducts 60 measurements.

**Table 4:** Exemplary plan for a method validation conventional interlaboratory study.

| Lab | Instrument performance | Class A | Class B |
| --- | --- | --- | --- |
| Lab 1 | superior instrument | 1-30 | 1-30 |
| Lab 2 | superior instrument | 31-60 | 31-60 |
| Lab 3 | average instrument | 1-15 and 31-45 | 1-15 and 31-45 |
| Lab 4 | average instrument | 16-30 and 46-60 | 16-30 and 46-60 |

***Step 2:*** Of the 60+60 samples, 3+3 samples are selected for which the correct classification proved to be particularly difficult in Step 1, with decision scores close to the threshold. A further 12+12 samples are chosen randomly. A total of 15+15 samples are finally tested in a conventional interlaboratory comparison with six laboratories without replicates. Therefore, each of the six laboratories conducts 30 analyses.

***Step 3:*** In the statistical evaluation, the results of both Step 1 and Step 2 are considered. First, the samples are tested for variance homogeneity. If it turns out that substantial variances occur for the 3+3 challenging samples, these samples can be considered separately. Then the laboratories are checked for variance homogeneity. Suppose variance is homogeneous across laboratories. In that case, an analysis of reproducibility, matrix variability, and error probabilities can be performed across laboratories and samples (taking into account inter-laboratory and inter-sample variations). However, if the assumption of variance homogeneity is violated for the laboratories or a standardized evaluation is not possible, e.g., by taking QC data into account, the data must be evaluated separately for each laboratory. This means that although it is possible to standardize the method, validation can only be performed by taking into account the individual laboratory.

### 3.3 Method validation interlaboratory study (factorial)

Ideally, the NTM must be performed several times for each reference sample to be able to detect analytical effects. Factors that could influence the measurement due to the variation of storage and sample preparation (e.g., water content, temperature or kind of extraction, hydrolysis, digestion, or proteolysis) should be specifically varied, if necessary. Here, the use of factorial experiments is recommended. Within a factorial design study, the characterization of the standard sample should represent not only the variability and bandwidth of the population (total production) but also the bandwidth of what can influence the measured spectra during storage, sample preparation, analysis, and evaluation. Relevant information on these parameters and information on the analysis itself (laboratory, instrument, reagents, processor, time of analysis, etc.) should be kept available if possible, at least as far as data protection requirements do not prohibit it.

The total number of tests to be performed and the number of laboratories involved can be reduced if the interlaboratory comparison is performed according to a factorial design. Only four laboratories are needed, but the performance of the laboratories should be similar, preferably using comparable instruments.

***Step 1:*** Four laboratories perform measurements for a total of 60 representative samples of population A and 60 representative samples of population B according to a standardized procedure with uniform sample preparation. Each sample is analyzed by one lab only, and each of the 120 samples is analyzed once. Each laboratory conducts 15+15 analyses.

***Step 2:*** Then, eight samples are randomly selected from each of the two populations (A & B) and analyzed without repetition according to an exemplary design described in Table 5. Each of the four laboratories conducts 32 analyses. The table illustrates a design with four factors, time, storage time, operator, and sample preparation. The analysis is done across laboratories.

**Table 5:** Factorial design plan for an inter-laboratory method validation study. Ones and zeros represent two-factor levels for the operator (e.g., operator E and F), sample storage time (long and short), and cartridge (e.g., purchased from company G and H, or belong to different lots.).

| Sample | Population | Day | Operator | Storage time | Cartridge | Lab 1 | Lab 2 | Lab 3 | Lab 4 |
| --- | --- | --- | --- | --- | --- | --- | --- | --- | --- |
| 1 | B | 1 | 1 | 1 | 1 |  |  |  |  |
| 1 | B | 1 | 2 | 2 | 2 |  |  |  |  |
| 2 | B | 1 | 1 | 2 | 1 |  |  |  |  |
| 2 | B | 1 | 2 | 1 | 2 |  |  |  |  |
| 3 | B | 1 | 2 | 1 | 2 |  |  |  |  |
| 3 | B | 1 | 1 | 2 | 1 |  |  |  |  |
| 4 | B | 1 | 2 | 1 | 2 |  |  |  |  |
| 4 | B | 1 | 1 | 2 | 1 |  |  |  |  |
| 5 | B | 2 | 1 | 1 | 1 |  |  |  |  |
| 5 | B | 2 | 2 | 2 | 2 |  |  |  |  |
| 6 | B | 2 | 1 | 2 | 1 |  |  |  |  |
| 6 | B | 2 | 2 | 1 | 2 |  |  |  |  |
| 7 | B | 2 | 2 | 1 | 2 |  |  |  |  |
| 7 | B | 2 | 1 | 2 | 1 |  |  |  |  |
| 8 | B | 2 | 2 | 1 | 2 |  |  |  |  |
| 8 | B | 2 | 1 | 2 | 1 |  |  |  |  |
| 9 | A | 1 | 1 | 1 | 1 |  |  |  |  |
| 9 | A | 1 | 2 | 2 | 2 |  |  |  |  |
| 10 | A | 1 | 1 | 2 | 1 |  |  |  |  |
| 10 | A | 1 | 2 | 1 | 2 |  |  |  |  |
| 11 | A | 1 | 2 | 1 | 2 |  |  |  |  |
| 11 | A | 1 | 1 | 2 | 1 |  |  |  |  |
| 12 | A | 1 | 2 | 1 | 2 |  |  |  |  |
| 12 | A | 1 | 1 | 2 | 1 |  |  |  |  |
| 13 | A | 2 | 1 | 1 | 1 |  |  |  |  |
| 13 | A | 2 | 2 | 2 | 2 |  |  |  |  |
| 14 | A | 2 | 1 | 2 | 1 |  |  |  |  |
| 14 | A | 2 | 2 | 1 | 2 |  |  |  |  |
| 15 | A | 2 | 2 | 1 | 2 |  |  |  |  |
| 15 | A | 2 | 1 | 2 | 1 |  |  |  |  |
| 16 | A | 2 | 2 | 1 | 2 |  |  |  |  |
| 16 | A | 2 | 1 | 2 | 1 |  |  |  |  |

The factorial approach’s particular advantage is that it should allow standardization for at least one type or generation of devices because the number of laboratories is now only half as large. In addition, the total number of required analyses is considerably lower at 248 compared to the conventional approach with 420 analyses. However, a disadvantage of the factorial approach is that it is not possible to evaluate the samples separately for each laboratory. This would require more samples.

### 3.4 Requirements for data management

Turning now to considerations and requirements for measurement data management, the following aspects are suggested. The participating laboratories should provide information on the storage of the samples (storage duration, storage temperature) and, if necessary, information on the condition of the samples (e.g., moisture content). Also important is information on the instruments, software used, reagents, and personnel (information on experience with the method), and the time period of the measurements. Furthermore, it is important to provide the measured data in a uniform form. For this purpose, the participants should be contacted in advance. This also includes making the spectra or sequence available, typically for reference material in a standardized form according to a uniform procedure. Suppose it turns out that the spectra or sequence shows a difference, e.g., a difference in instrument resolution. In that case, the laboratories should not modify the acquired data (e.g., coarsen the resolution); instead, leave this to the interlaboratory comparison study organizer.

Besides, additional metadata of the investigated samples should be provided, where possible. It is also beneficial for documentation purposes if the instrument’s information and the respective analytical data acquisition procedure are provided. It would be immensely useful if the laboratories also provided the spectra or sequences obtained from regular quality control analyses. Each spectrum or sequence submitted by the lab must accompany a laboratory ID to carry out laboratory-specific evaluations over a more extended period. If the QC samples are comparable, these data can be used to determine the temporal fluctuations that can be expected within the laboratories and whether these fluctuations are comparable from laboratory to laboratory. Finally, it is recommended that after completion of the interlaboratory comparison study, all measured data should be available centrally for all samples in a uniform manner (e.g., same data resolution) in order to give an impression of the variability between the spectra or sequences of different laboratories on the one hand, but also of the variability within the standard sample due to biological and process variations on the other hand.

To ensure comparability of the results in the interlaboratory comparison, the metadata of the samples distributed in the interlaboratory comparison must be determined or checked centrally. This also includes the verification of homogeneity and stability on the basis of selected peaks or sequences in a laboratory. Given the copious amounts of measurement data in the form of fingerprints and a multitude of algorithms to parse them, the data evaluation in itself can add to the variability of the measured result. Hence, a distinction between wet lab (involving all steps on the lab bench) and dry lab (involving data evaluation steps) can be made. There can be significant merit in evaluating the wet lab and dry lab precision parameters separately. Additionally, the calculation method parameters used in the dry lab steps must be made available centrally to enable the laboratories to evaluate their measured data. Independent of this, the evaluation of the data is recommended to be done centrally to determine whether differences between the evaluation by the participants and the central evaluation can be identified.

Even after the interlaboratory comparison study has concluded, the database should be updated regularly to provide current reference samples in the database. These newly inserted reference samples are subject to the same metadata requirements as the external quality assurance service samples. This can allow for long-term tracing and tracking of the sample characteristics – for example, change in climatic or agricultural patterns does not lead to a gradual change in the proteome of plant origin products. Alternatively, modifications to food production or packaging steps can lead to a change in the fingerprints. In these circumstances, the NTM would be rendered fruitless, as the procedure would no longer correctly distinguish between authentic and non-authentic samples.

## 4. Statistical evaluation

In the following, it is assumed that there is a quantitative decision score “z”, based on measurement *X*_*i*_. As per Equation 2, then *y*(*X*_*i*_) = 1 represents the case where *z*(*X*_*i*_) > *θ*, and *y*(*X*_*i*_) = 0 where *z*(*X*_*i*_) ≤ *θ*. Here, *θ* denotes the threshold value for the decision between A and B. z(*X*_*i*_) reflects the decision strength, showing the confidence if the sample belongs to population A (lower score) or B (higher score).

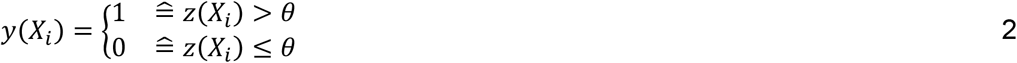

It is noteworthy to point out here that the decision score Z may not be metrologically traceable. It could be the result of an unknown or proprietary algorithm, such as an instrument manufacturer’s score provided through its software.

Let *Z*_*ijk*_ denote the decision score for sample *i*, laboratory *j* in the repeated measurement *k*. The basic model for classification based on quantitative scores is as follows:

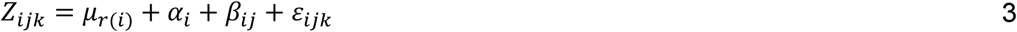

Here,

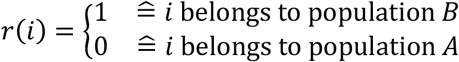

*μ*_1_ = mean of population *B*

*μ*_0_ = mean of population *A*

*α*_*i*_ = (random) deviation of sample *i*

*β*_*ij*_ = (random) deviation of the results of laboratory *j* for sample *i*

*ε*_*ijk*_ = random deviation of result *k* of laboratory *j* for sample *i* We also use the notation

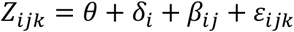

where *δ*_*i*_ = *μ*_*r*(*i*)_ + *α*_*i*_ − *θ. δ*_*i*_represents the difference between the threshold and the mean value of sample *i* across all laboratories.

Under assumptions of homogeneity of variances of laboratory deviations and random deviations within the population *A* and within the population *B*, the following variances can be estimated by means of a two-way random effect analysis of variance (ANOVA) –

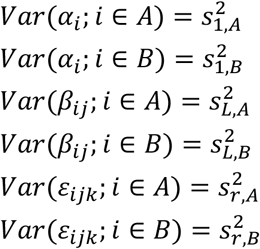

Based on the above variance components, we calculate the classification variances according to Equations 4 and 5. The smaller the classification variances, the better the classification power.

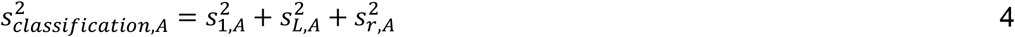

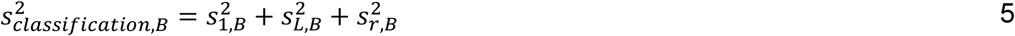

These classification variances consist of two components, the sample variances 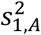 and 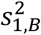 in populations A and B, and the reproducibility variances 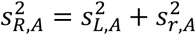 and 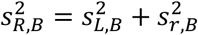 calculated as averages across A and B, respectively.

If the decision score can be considered normally distributed, then the sensitivity and specificity can be estimated using Equations 6 and 7. Here Φ denotes the cumulative distribution function of the standard normal distribution.

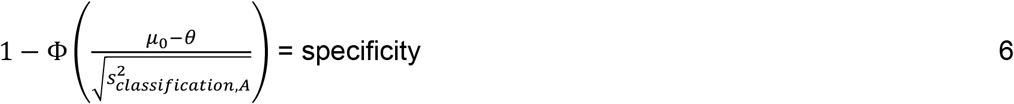

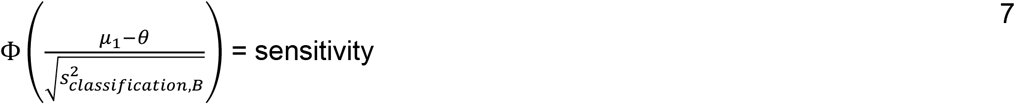

Here, Φ(.) denotes the cumulative distribution function of the standardized normal distribution. The equations are only an approximation since it cannot be assumed that the distribution of the decision scores corresponds exactly to the normal distribution. Noteworthy that the variances used are empirically determined values that are subject to random fluctuations.The model presented here allows for calculating error probabilities for the underlying populations A and B under the assumption that the sample was randomly selected. However, the informative value of these error probabilities is limited if the error probability is highly dependent on the selected sample itself. This corresponds to the case where the variances for the sample effect, 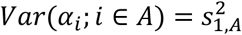 and 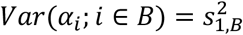, turn out to be relatively high, i.e., if the sample variances 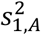 and 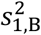 are higher than the respective reproducibility variances 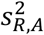 and 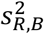 (see Figure 3) If the sample, i, is already known, calculate the sensitivity or specificity for this sample for the fixed, non-random value *δ*_*i*_

**Figure 3.**
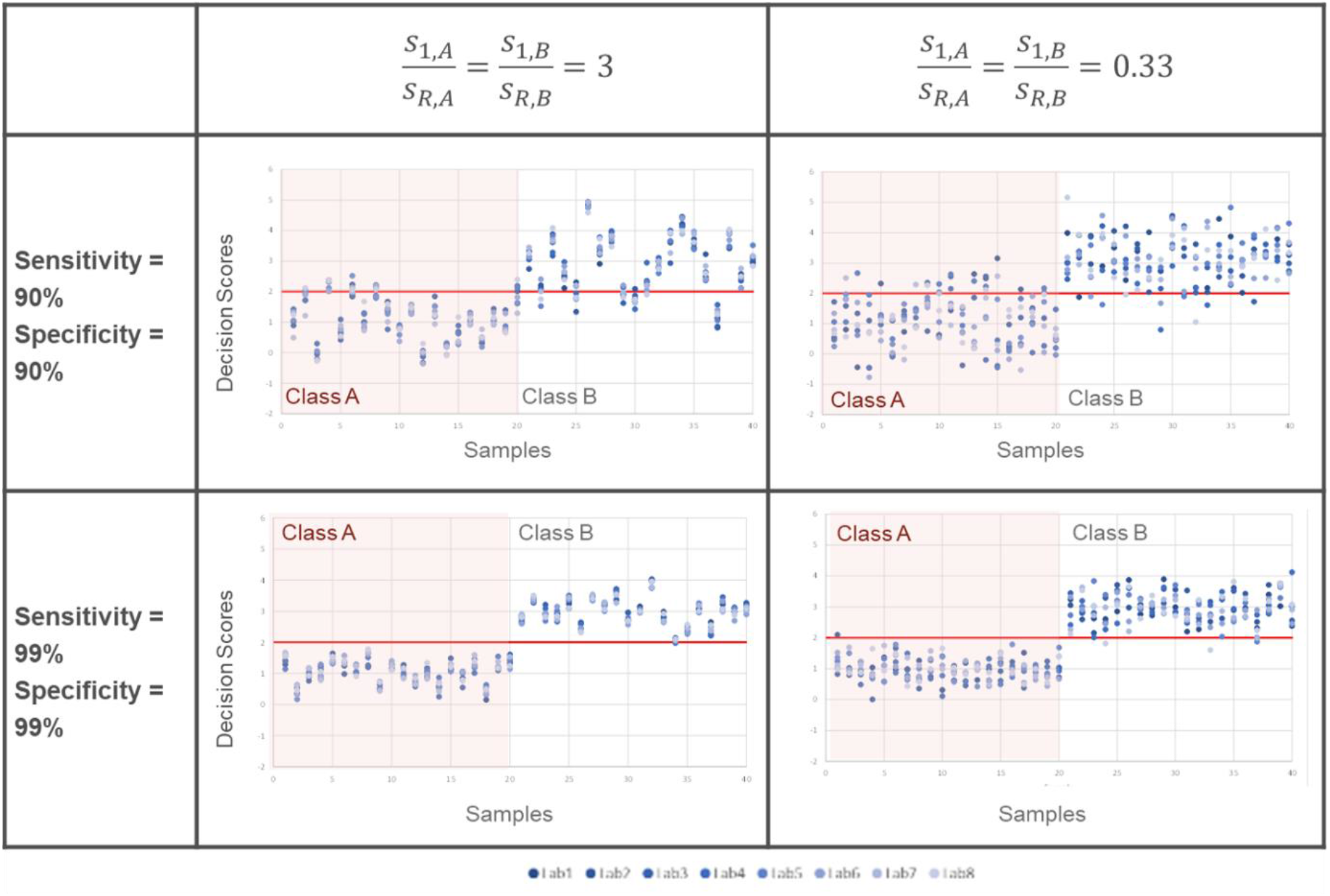
Exemplary distribution of decision scores for 20+20 samples belonging to class A and B, under different sensitivity and specificity conditions. Different shades of blue dots show the measurement values for different laboratories (n=8). A decision threshold of 2 is considered.

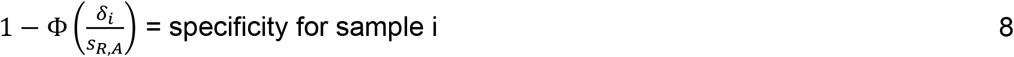

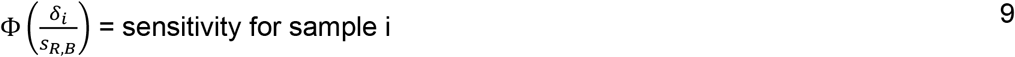

The variance components calculated here can also be used to determine the samples’ respective sensitivity or specificity and, consequently, evaluate the results’ measurement uncertainty based on the decision score. Finally, these variance components can also be used to capture, on the one hand, the effects of chemical-analytical or biochemical analysis (wet lab) and, on the other hand, the effects resulting from different forms of data analysis (dry lab).

Consider a simulation of an interlaboratory method validation study. Although 60 samples for each class A and B were recommended in the previous sections, Figure 3 shows an excerpt of the results from 8 laboratories for 20 samples per class. On measuring all the samples using an NTM, a decision score is obtained. A threshold for a decision threshold as determined in the method development stage is shown by the red horizontal line for a score of 2. Each of the eight labs reports a decision score (as shown by the different shades of blue dots).

Figures 3 and 4 illustrate two different scenarios in the two rows - (i) when the average sensitivity and specificity are high 90%, and (ii) when both the average sensitivity and specificity are very high, i.e., 99%. These scenarios can be illustrative of a good performing method and not so good method. For the cases in figures in the left panels, the variation between samples is large. But the variation over different laboratories is relatively small. Variation in samples can be due to differences in processing, taxonomy or species, geographical origin, harvest time/conditions, etc. What stands out is the result for certain samples, e.g., #37, which has a decision score below the threshold of 2 as measured by all the labs. Besides, the outcome of samples shows large disparities when measured by different laboratories (e.g., #2, #4, #6, #8, #20, #22, etc.). Such ‘challenging’ samples can show larger variation around the decision threshold. In another case, figures in the right panels show a small variation between samples but a large reproducibility variation. Thus here, the laboratory variance is of greater relevance in the validation study. This can be due to several factors such as a difference in data analysis pipeline or database, extraction techniques, instrumentation, or operator experience. Furthermore, such a situation can also arise due to a difficult matrix for the food sample, which leads to large variations in how the labs perform the extraction process.

**Figure 4.**
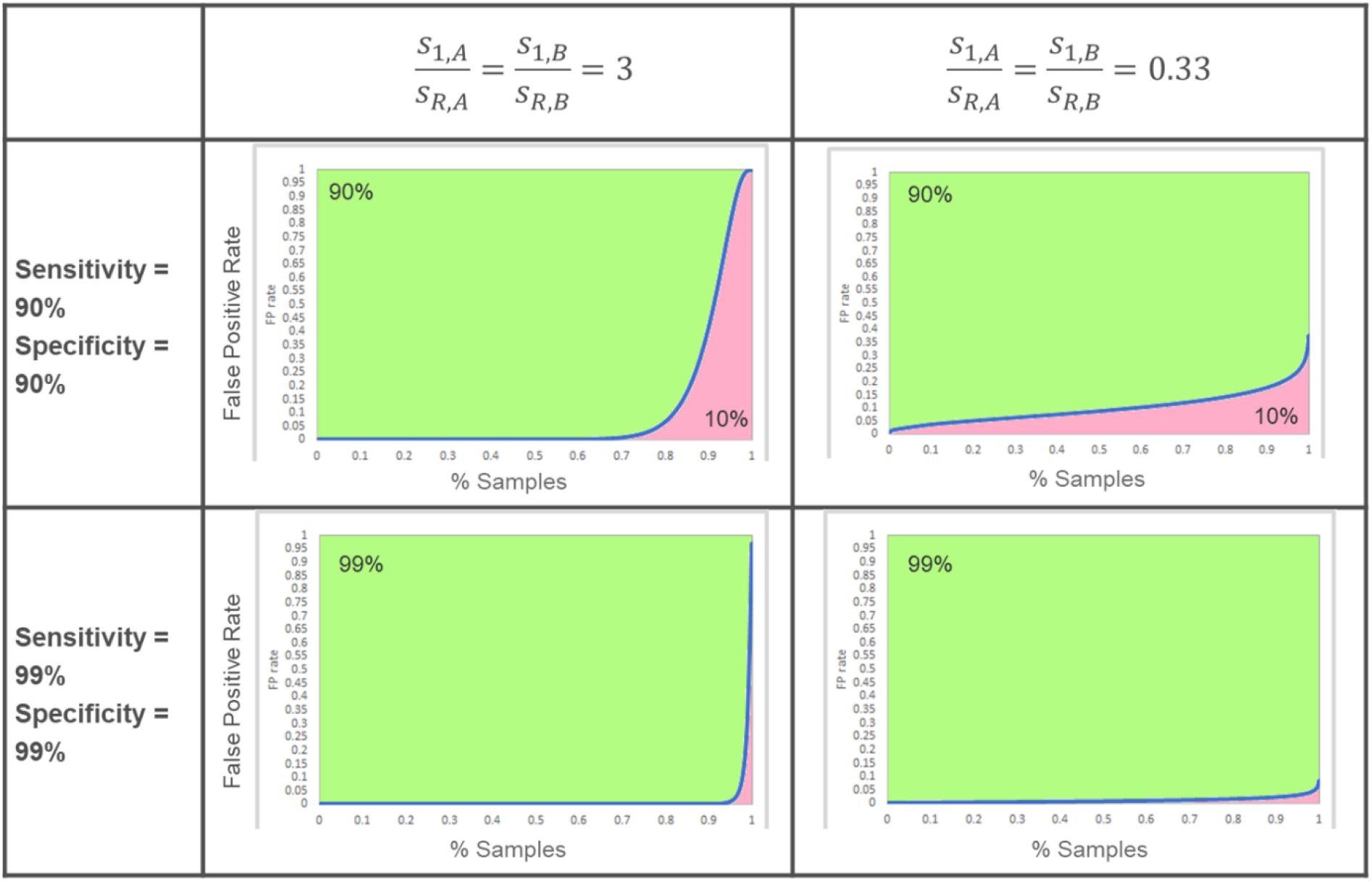
False positive rate changing with the theoretical percentage of all authentic samples for different scenarios.

Even though such simulated scenarios only form a small subset of how method validation studies can turn out, what is clear from them is that despite the methods having the same average false-negative rate and false-positive rates, careful considerations should be made to make unequivocal claims about the performance of the method.

To further understand how the sample variation and reproducibility variation – together influence the outcome of the validation study, Figure 4 shows the conditional false positive (FP rates), which is (1-specificity) for the four different scenarios. It can be seen that if the specificity = 90% and 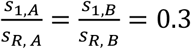 (upper left chart), then for 80% of all authentic samples (population, the FP rate is below 0.07, and for 90% of all authentic samples, the FP rate is below 0.45. Conversely, it can be concluded that for 20% of all authentic samples, the FP rate is above 0.07, and for 10%, even above 0.45. Here, the variability between samples is dominant. If the analytical variability is dominant (upper right chart), the variability of the FP rate is less pronounced. It should be noted that in both scenarios, the mean FP rate across all samples is 0.1, which corresponds to a specificity of 90%.

At a specificity of 99%, the FP rate is on average 0.01. With a dominating sample standard deviation (lower left figure), false-positive results are concentrated on approximately 5% of all samples. For 95% of all authentic samples (population A), the FP rate is below 0.01, and for 99% of all authentic samples, the FP rate is below 0.37. Conversely, it can be concluded that for 5% of all authentic samples, the FP rate is above 0.01, and for 1% even above 0.37. With dominating analytical variability (lower right figure), the variability of the FP rate is less pronounced.

The scenarios described above illustrate the role of challenging samples in validation studies. For the design of validation studies, this raises the question of which criteria can be used to decide whether a sample is challenging. Often this may be known, for example, when particularly complex matrices or sample mixtures are involved. In such a case, it may be possible to consider “normal” and challenging matrices separately. However, if it is not known which samples are challenging, the task arises in the context of a validation study to evaluate the ratios 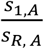 and 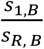. If it is possible to identify challenging samples before the NTM method is applied, more specific discrimination strategies could be applied in order to minimize false positive and false negative rates. Therefore, the experimental design of the validation study also depends on whether challenging samples are identifiable as such before the actual analysis.

## 5. Conclusion

For NTMs to achieve wide acceptance and be used in court, sufficient accuracy of the results must be ensured. This report describes a validation strategy for binary classification type NTMs. Besides, it highlights the need for considerations of different sources of variability, which should be taken into account. The rationale for the required number of samples to be measured can be of significant interest to method developers, users, and regulatory organizations. Furthermore, the report highlights the distinction between a wet lab (analytical measurement on the lab bench) and a dry lab (downstream data evaluation) components in any NTM, which can add to the variation in the measured values. Through a simulation study, it is demonstrated that even with the same performance characteristics (sensitivity and specificity), on a closer appraisal of the method, the false positive rate within the population of authentic samples can vary. Furthermore, it is apparent that the question of when is a sample challenging – is an additional important consideration. It also follows that not all NTMs should be evaluated using the same experimental design but that special attention should be paid to challenging but authentic samples. Altogether, this report contributes to the groundwork being laid for developing and harmonizing validation schemes for NTMs.

